# How Array Design affects SNP Ascertainment Bias

**DOI:** 10.1101/833541

**Authors:** Johannes Geibel, Christian Reimer, Steffen Weigend, Annett Weigend, Torsten Pook, Henner Simianer

## Abstract

Single nucleotide polymorphisms (SNPs), genotyped with SNP arrays, have become a widely used marker type in population genetic analyses over the last 10 years. However, compared to whole genome re-sequencing data, arrays are known to lack a substantial proportion of globally rare variants and tend to be biased towards variants present in populations involved in the development process of the respective array. This affects population genetic estimators and is known as SNP ascertainment bias. We investigated factors contributing to ascertainment bias in array development by redesigning the Axiom™ Genome-Wide Chicken Array *in silico* and evaluating changes in allele frequency spectra and heterozygosity estimates in a stepwise manner. A sequential reduction of rare alleles during the development process was shown with main influencing factors being the identification of SNPs in a limited set of populations and a within-population selection of common SNPs when aiming for equidistant spacing. These effects were shown to be less severe with a larger discovery panel. Additionally, a generally massive overestimation of expected heterozygosity for the ascertained SNP sets was shown. This overestimation was 24% higher for populations involved in the discovery process than not involved populations in case of the original array. The same was observed after the SNP discovery step in the redesign. However, an unequal contribution of populations during the SNP selection can mask this effect but also adds uncertainty. Finally, we make suggestions for the design of specialized arrays for large scale projects where whole genome re-sequencing techniques are still too expensive.

## Introduction

Starting in the first decade of this century, the possibility of cost effectively genotyping high numbers of Single Nucleotide Polymorphisms (SNP) for many individuals in parallel via SNP arrays led to an increased usage of them for population genetic analyses in humans [1, 2], model species [3, 4], plants [5, 6] and livestock [7–13].

Various SNP arrays exist for humans [14], plants [15, 16] and all major livestock species [17–23]. SNP numbers within these arrays range from 10 k SNPs [20] over a range from 40 to 60 k [16, 17, 19, 22] up to 600 k [15, 21]. The design process of every array has an initial step of SNP discovery in common, where SNPs are identified from existing databases and/or from a small set of sequenced individuals. SNPs are then selected based on different quality criteria like minor allele frequency (MAF) thresholds and platform specific design scores [24]. Additional criteria like equidistant spacing over the genome [21], overrepresentation of some areas [20] or increased overrepresentation of high MAF SNPs [17] are applied dependent on the design intentions. In the end, draft arrays are validated either on the set of populations used for the SNP discovery itself [18] and/or on a broad set of individuals from different populations [21, 24].

In contrast to whole genome re-sequencing (WGS) data, SNP arrays often show a clear underrepresentation of SNP with extreme allele frequencies [25]. As population genetic statistics are mostly based on estimates of allele frequencies, this context leads to biased population genetic estimators [25, 26] and is known as SNP ascertainment bias.

The absence of rare alleles is mainly driven by two factors in the array design process where SNPs are selected (ascertained) based on different requirements and decisions [27]. The first factor is a relatively small panel of individuals being used for discovery of SNPs, leading to a large proportion of globally rare variants not being selected, since they appear monomorphic in the discovery panel [26, 28]. The second factor is the across population use of arrays. Arrays are developed based on the variation within the discovery panel, thus missing variation present in distantly related individuals or populations [25, 27]. This second source of bias was shown to be of relatively high importance for livestock studies, where arrays are usually developed for large commercial breeds and later used to genotype diverse sets of local breeds all over the world [29, 30].

Besides different strategies to minimize the impact of ascertainment bias [30, 31], there are some attempts to correct the allele frequency spectrum via Bayesian methods [25, 28, 32]. However, those corrections highly rely on detailed statistical assumptions of the ascertainment process [33, 34] and are currently only tested for corrections of the first source of ascertainment bias, the small discovery panel [25, 28, 32]. In practice, detailed information on the design process is limited [34] and the complexity of the processes makes statistical models for the corrections inaccurate.

Agricultural species such as chickens often show a complex domestication history, and therefore allow for few prior assumptions on ascertainment bias. Domestic chickens are assumed to originate from red jungle fowl (*Gallus gallus*) ancestors in Southeast Asia [35, 36], represented by the five subspecies *G. g. gallus, G. g. spadiceus, G. g. murghi, G. g. bankiva* and *G. g. jabouillei* [37]. Additionally, some hybridization events with other *Gallus* species (e.g. grey jungle fowl; *Gallus sonneratii*) have been suggested [36, 38]. The diversity of today’s local breeds of chickens in Europe originates from chickens that reached the continent about 3000 years ago via a northern and a southern route, followed by selection and crossing with Asian chicken breeds introduced in the 19th century [37]. While commercial white layers were derived solely by intensive directional selection of a single breed, the White Leghorn, commercial brown layers are derived from a broader genetic basis (e.g. Rhode Island Red, New Hampshire, Barred Plymouth Rock). Commercial broilers are derived by cross-breeding of paternal lines (e.g. White Cornish) with maternal lines which descend from a comparable basis as brown layers (e.g. White Plymouth Rock) [39]. For more detailed information on chicken ancestry we refer to Qanbari *et al*. [36] and for a comprehensive overview on diversity and population structure of domesticated chickens to Malomane *et al*. [13].

Given the complexity of modern array design processes and the chicken population structure, this study aims at highlighting the mechanisms which promote the bias by illustrating the effects of the different steps of the array design process on ascertainment bias, using real data in a typical setting from livestock sciences. For this purpose, the design process of the Axiom™ Genome-Wide Chicken Array [21] was simulated in a set of diverse chicken WGS data. Allele frequency spectra as well as expected heterozygosity (H_exp_) were compared to the WGS data and the SNPs of the Axiom™ Genome-Wide Chicken Array. Finally, some recommendations are made to design an array for monitoring genetic diversity.

## Material and methods

### Populations and sequencing

The analysis is based on WGS data of a diverse set of 46 commercial, non-commercial and wild chicken populations, sampled within the framework of the projects AVIANDIV (www.aviandiv.fli.de) and SYNBREED (www.synbreed.tum.de). Commercial brown (BL) and white layer (WL) populations consist of 25 individually re-sequenced animals each, while the two commercial broiler lines (BR1 and BR2) include 20 individually sequenced animals each. For 41 populations, pooled DNA from 9 - 11 animals per population was sequenced, while *Gallus varius* (green jungle fowl; GV) samples of only two animals were sequenced as a pool. More detailed information about the samples can be found in **S1 File** and two previously published papers, from Malomane *et al*. [30] and Qanbari *et al*. [40]. Coverage was between 7X and 10X for the individual sequences, while DNA pools were sequenced with 15X to 70X coverage. Sequencing was conducted on Illumina HiSeq machines at the Helmholtz Zentrum, German Research Center for Environmental Health in Munich, Germany.

### Raw data preparation and SNP calling

Sequences were aligned to the reference genome Gallus gallus 5.0 [41, 42] and the SNP calling was conducted according to GATK Best Practices guidelines [43, 44]. BWA-MEM 0.7.12 [45] was used for the alignment step, duplicates were marked using Picard Tools 2.0.1 [46] MarkDuplicatesWithMateCigar and base qualities were recalibrated with GATK 3.7 [47] BaseQualityRecalibrator. The set of known SNPs, necessary for base quality score recalibration, was downloaded from ENSEMBL release 87 [48]. SNPs were called for all samples separately using the GATK 3.7 HaplotypeCaller and later on simultaneously genotyped across samples with GATK 3.7 GenotypeGVCFs. Due to computational limitations, a ploidy of two instead of the higher true ploidy of the pooled sequences had to be assumed, which led to calling slightly less rare alleles. However, effects of this limitation are negligible **(S2 File; S1 Fig)**.

SNP filtering was conducted using GATK 3.7 VariantRecalibrator, which filtered the called SNPs by a machine learning approach (use of a Gaussian mixture model), which uses both, a set of previous known (low confidence needed) and a set of highly reliable (assumed to be true) variants as training resources [49]. The source for known SNPs (prior 2) provided to VariantRecalibrator was again ENSEMBL (release 87) and the SNPs of the Axiom™ Genome-Wide Chicken Array were defined as true training set (prior 15). The algorithm was trained on the quality parameters DP, QD, FS, SOR, MQ and MQRankSum. Filters were set to recover 99 % of the training SNPs in the filtered set, which resulted in a Transition/Transversion ratio of 2.52 for known SNPs, and a Transition/Transversion ratio of 2.26 for novel SNPs. Only biallelic autosomal SNPs were used in all further analyses.

### Identification of the ancestral allele

Ancestral alleles were defined using allele frequency information from the three wild populations *Gallus gallus gallus* (GG), *Gallus gallus spadiceus* (GS) and *Gallus varius* (GV) by an approach comparable to 50 [50]. It was assumed that the *Gallus gallus* and *Galllus varius* species emerged from a common ancestor and *Gallus gallus* later split into *Gallus gallus gallus* and *Gallus gallus spadiceus* subspecies. Additionally, assuming neutral molecular evolution [51], the ancestral allele was most likely the major allele within those three populations, when weighting the allele frequency of *Gallus varius* twice. This procedure assigned the ancestral status to the reference allele for 86 % of the SNPs and to the alternative allele for 14 % of the SNPs. The change in the allele frequency spectrum was only relevant for the interval from 0.95 – 1.00, which was reduced by 111’851 SNPs (0.39 % of all SNPs) when switching from alternative to derived allele frequency **(S2 File; S2 Fig)**.

### Reference Sets

Three different reference sets were defined as follows: the **unfiltered WGS** SNPs (28.5 M SNPs), SNPs filtered using GATK 3.7 [47] VariantRecalibrator (20.9 M SNPs;filtered WGS) and **array SNPs** (540 k SNPs), which are the intersection of the unfiltered SNPs and the SNPs of the Axiom™ Genome-Wide Chicken Array. The separate use of unfiltered and filtered WGS SNPs was done to assess the effect of filtering (especially the use of an ascertained SNP set as the true set) on ascertainment bias.

### Redesigning the SNP Array

The populations were divided into four groups for the array design:

1. Discovery populations (8)
2. Validation populations (19)
3. Application populations (18)
4. Outgroup (1)

For SNP discovery firstly the 4 commercial lines (commercial white layers, WL; commercial brown layers, BL and the two commercial broiler lines, BR1 and BR2) were used. The set was then extended by additionally selecting those populations that were closest related to each of the commercial populations based on pairwise Nei’s standard genetic distance [52] As the two broiler populations were closest related **(Fig 1**), the next two closest populations were chosen. This resulted in the inclusion of White Leghorn (LE), Rhode Island Red (RI), Marans (MR) and Rumpless Araucana (AR). Note that these populations are closely related to the populations used as discovery populations for the development of the Axiom™ Genome-Wide Chicken Array [21] with exception to some inbred lines from the Roslin Institute in Edinburgh we do not know the genetic origin of. The discovery set used for the original array [21] additionally consisted of more animals than ours. Further, the discovery set had to be split into broilers (BR1, BR2, MR, AR) and layers (WL, LE, BL, RI) for the equal spacing step. From the remaining populations, 19 were randomly chosen for SNP validation (validation populations), 18 populations (which were not included in the array development) were used as showcase for an application of the array (application populations), and *Gallus varius* as a different species was defined as outgroup. The interested reader can find all underlying pairwise Nei’s standard genetic distances [52] in **S3 File** and additionally pairwise F_ST_ values [53] in **S4 File**.

**Fig 1.**
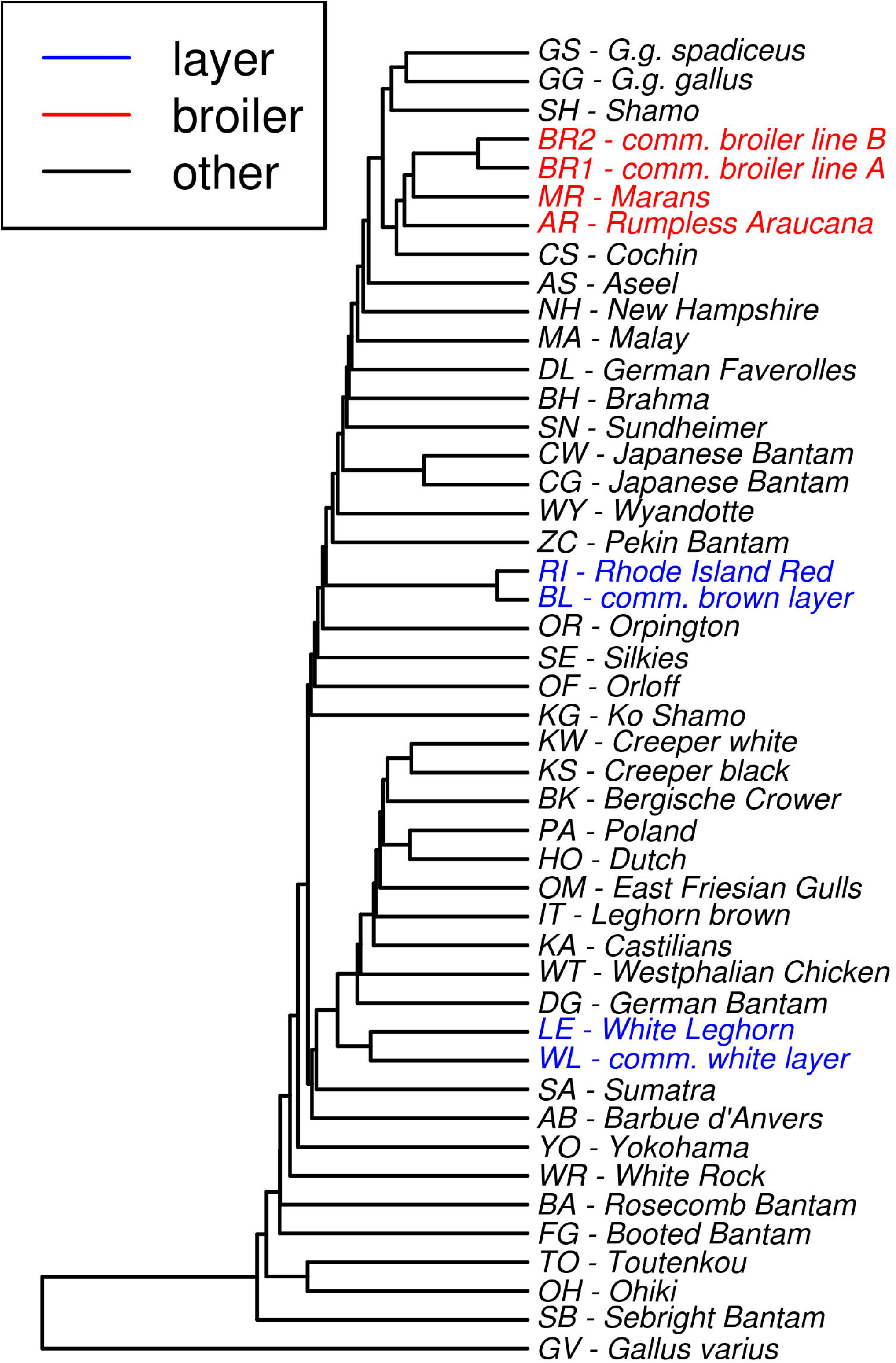
UPGMA tree based on pairwise Nei’s standard genetic distances. The tree was calculated from the filtered WGS SNPs. Populations defined as layers or broilers, which form in total the discovery set for the array design close to the original array, are highlighted. The plot was produced using the R package ape [54]. Note that the plot is only supposed to reveal close clustering chicken populations and cannot be interpreted in depth as chickens show a rich history of hybridization events. The interested reader can find all underlying pairwise Nei’s standard genetic distances [52] in **S3 File** and additionally pairwise FST values [53] in **S4 File**.

Based on the unfiltered SNP set, the sampling of the SNPs for an approximately 600 k sized array was remodeled *in silico* in five consecutive steps according to the design process of the original array which was described by Kranis *et al*. [21], starting from the unfiltered SNP set:

1. **SNP discovery → 10.9 M SNPs** Discovery of SNPs fulfilling basic criteria (quality ≥ 60; MAF ≥ 0.05; coverage ≤ mean + three standard deviations) within the discovery populations.
2. **Cluster removal → 8.8 M SNPs** SNP clusters with less than 4 bp invariant sites at one side of a SNP (no matter of whether 3’ or 5’ direction) and less than 10 bp invariant sites at the other side of the SNP within the discovery populations were removed. This step is justified rather technically to enable probe binding, but could also lead to an overrepresentation of conserved regions compared to highly variable regions of the genome.
3. **Equal spacing → 2.1 M SNPs** Reduction of SNPs to achieve approximately equidistant spacing between variable SNPs within discovery populations based on genetic distances. The used algorithm was modeled according to Kranis *et al*. [21] and followed a two-step procedure. The first step was setting up an initial backbone of common SNPs (three sub steps). It started with selecting SNPs which segregated in all discovery populations (MAF within each population > 0) while requiring a minimal distance of 2 kb, resulting in about 8 k SNPs. This was complemented by a backbone of SNPs which segregated in all layer populations and another one of SNPs which segregated in all broiler populations. Note that Kranis *et al*. [21] additionally constructed a backbone from a group of inbred lines for which no comparable samples were available for this study. In the second step, the algorithm iterated over all single populations and filled in potential gaps between backbone SNPs which are variable within the according population. This was done by choosing the SNPs closest to equidistant positions within the gap while aiming for a predefined local target density of 667 segregating SNPs/cM (linkage map taken from [55]). See **S3 Fig** for the detailed contribution of additional SNPs from each sub step of the algorithm.
4. **SNP validation → 1.7 M SNPs** Removing SNPs (^~^ 20 %) which were not variable in at least 8 of the 19 validation populations. This step would in reality be done by genotyping with preliminary test arrays and therefore allows the use of a broader set of populations than the discovery step.
5. **Downsampling → 580 k SNPs** Downsampling of SNPs comparable to step 3, but without adding the broiler/ layer specific backbones and instead keeping all exonic SNPs (annotation using Ensembl VEP 89.7; 56). Additionally, the target density in broiler lines was set as three times the target density of the layer lines. The increased target density in broilers is intended to account for lower levels of linkage disequilibrium in these lines.

### Variation of the design process

The whole design process was repeated 50 times with populations being randomly assigned to be discovery, validation or application populations, while the *Gallus varius* population was always kept as the outgroup. In this process, the number of populations per group was the same as in the previous scenario.

To assess the impact of the number of discovery populations on the design process, the number of discovery populations was varied in additional runs from 4 to 40 randomly chosen populations (while assigning the remaining populations, except *Gallus varius*, to validation and application groups of equal size) with 20 random replicates for each number of discovery populations. In a last scenario, equal spacing was varied with respect to the target density (33 – 3333 SNPs/cM) with 20 independent population groupings for each target density, with or without the initial backbone. As the number of SNPs from the backbone was constant, the increase of the target density led to a higher number of SNPs chosen by the algorithm due to the equal spacing itself and hence the relative influence of the fixed number of common backbone SNPs decreased.

### Analyses of the results

Per-locus-allele frequencies for individually sequenced populations were estimated from genotypes, whereas the estimation for the sequenced DNA-pools was based on the allelic depth. Influences on the allele frequency spectra were examined by comparing density estimates of derived allele frequency spectra (unfolded frequency spectrum). Further H_exp_, the expected heterozygosity assuming Hardy Weinberg frequencies of the genotypes, for the different populations were used as summary statistics of the within population allele frequency spectra and calculated as in equation (1), where *p_ref;1_* denotes the frequency of the reference allele at locus *l* and *L* the total number of loci.

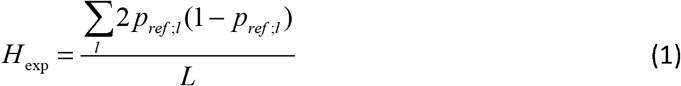

Deviations in the estimation of H_exp_ from the various SNP sets were quantified as differences between the H_exp_ calculated from the respective SNP set and the H_exp_ calculated from the filtered WGS SNPs relative to the H_exp_ from the filtered WGS SNPs, further called overestimation of H_exp_ (OHE; equation (2)). An OHE of zero means that the estimates are equal, while an OHE of one describes doubling of the unbiased estimate.

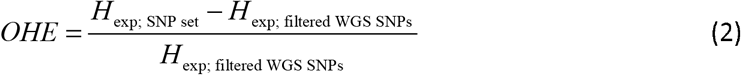

The effects of the population group assignments on the OHE of the random population assignments were evaluated by pairwise comparisons of least square means (LSMEANS; calculated with the R package emmeans [57, 58] by using Tukey correction for multiple pairwise contrasts) of the population groups. An underlying mixed linear model for the estimation of LSMEANS was fitted using the R package lme4 [59] as shown in equation (3), where the OHE depended on an overall mean *μ*, the fixed effect of the population group *popG_i_* (i can be discovery-, validation-, application- or outgroup), a random effect for the j^th^ repetition of random population grouping 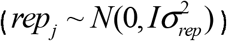 and a random error 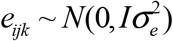. The procedure is comparable to simple pairwise comparisons of group means, the correction by the repetition only reduces the error variance and thus decreases the confidence intervals.

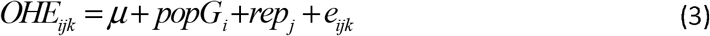

## Results

### Numbers of SNPs

The SNP calling identified 28.5 M biallelic autosomal SNPs from which 20.9 M SNPs passed GATK’s filtering procedure. 540 k SNPs from the unfiltered WGS SNP set are also mapped on the original Axiom™ Genome-Wide 580 k Chicken Array. The remodeling of the array according to the design process of the original array returned 10.9 M SNPs from the discovery step, which were reduced to approximately 580 k in steps as described. Numbers of identified SNPs for the additional runs differed depending on the populations and settings used and are listed in **S1 Table**. It has to be noted that the different sub steps of the equal spacing algorithm contributed with different amounts of SNPs **(S3 Fig)**. Especially the much higher contribution of SNPs which were segregating in all broiler populations compared to SNPs segregating in all layer populations in the first remodeling, which is due to closer relationships between the broiler populations and their generally higher heterozygosity, was remarkable. Additional information about the identified number of SNPs depending on the number of discovery populations and target density as well as information about the share of SNPs of different random runs can be found in **S4 – S6 Figs**.

### Underrepresentation of rare SNPs

A clear underrepresentation of rare SNPs in all ascertained SNP sets compared to WGS is evident from the allele frequency spectra **(Fig 2)**. Major changes in the allele frequency spectra during the array development process were observed after the SNP discovery step and the equal spacing step. The SNP discovery led to an underrepresentation of rare SNPs compared to sequence data, which was intensified by the equal spacing step **(Fig 2)** and resulted finally in a spectrum close to the spectrum of the original array. Randomly choosing populations as discovery populations confirmed the shape of the first remodeling, where the population groups were chosen according to the original array [21]. As major changes in the spectra mainly occurred after the SNP discovery and equal spacing, further results will concentrate on those steps.

**Fig 2.**
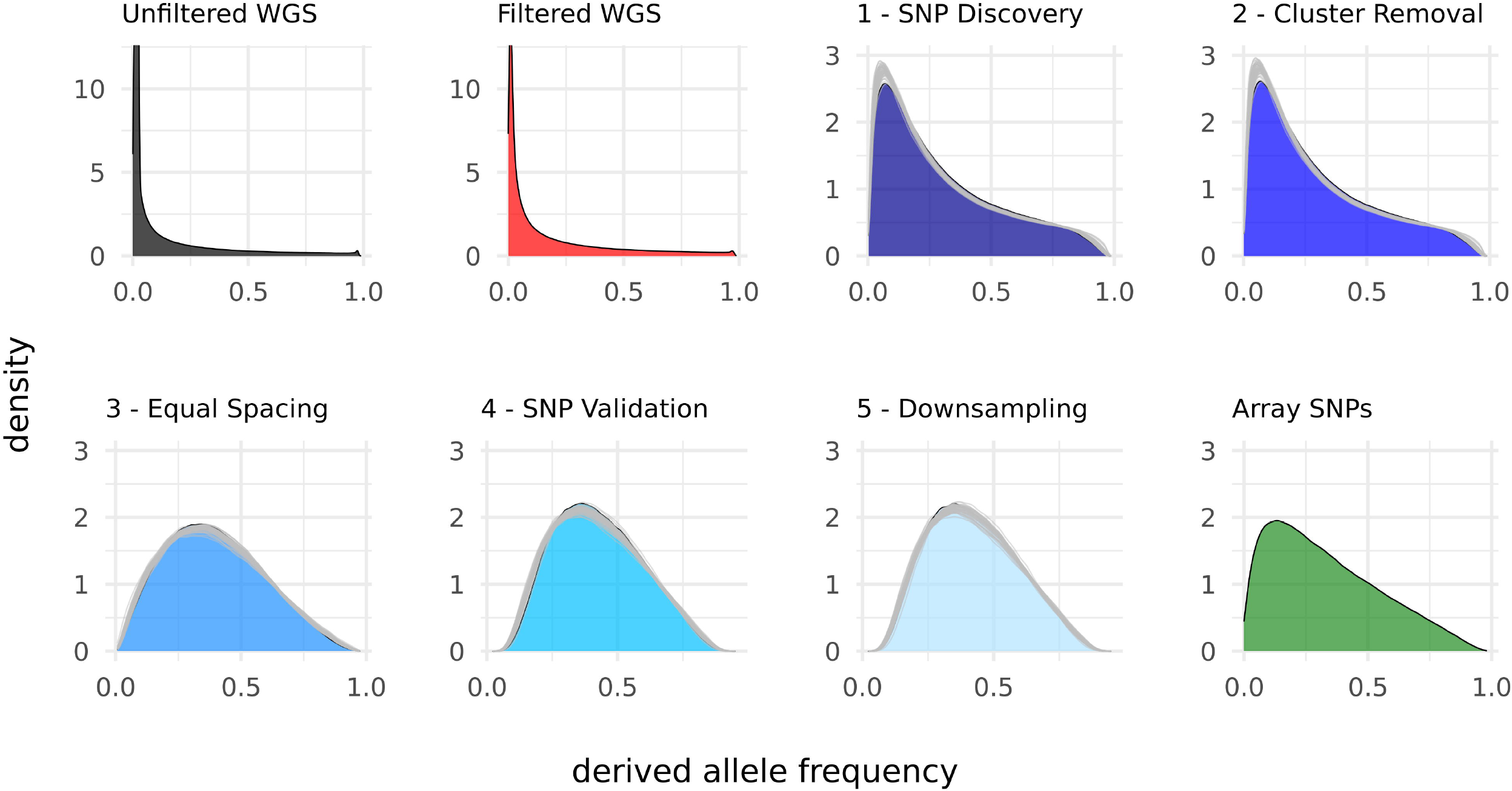
Derived allele frequency spectra for the different SNP sets. For the remodeled sets, areas show the modelling according to the original array [21] while grey lines represent the 50 random population groupings.

The allele frequency spectra **(Fig 3)** within discovery populations, compared to the spectra over all populations, clearly showed the cutoff from the MAF 0.05 filter. Furthermore, the allele frequency spectra of the discovery populations revealed a higher share of common SNPs than the overall spectra after equal spacing. In contrast, the spectra within validation- and application populations showed less pronounced peaks after the discovery step and the outgroup (*Gallus varius*) revealed fixation of most SNPs variable in the discovery populations.

**Fig 3.**
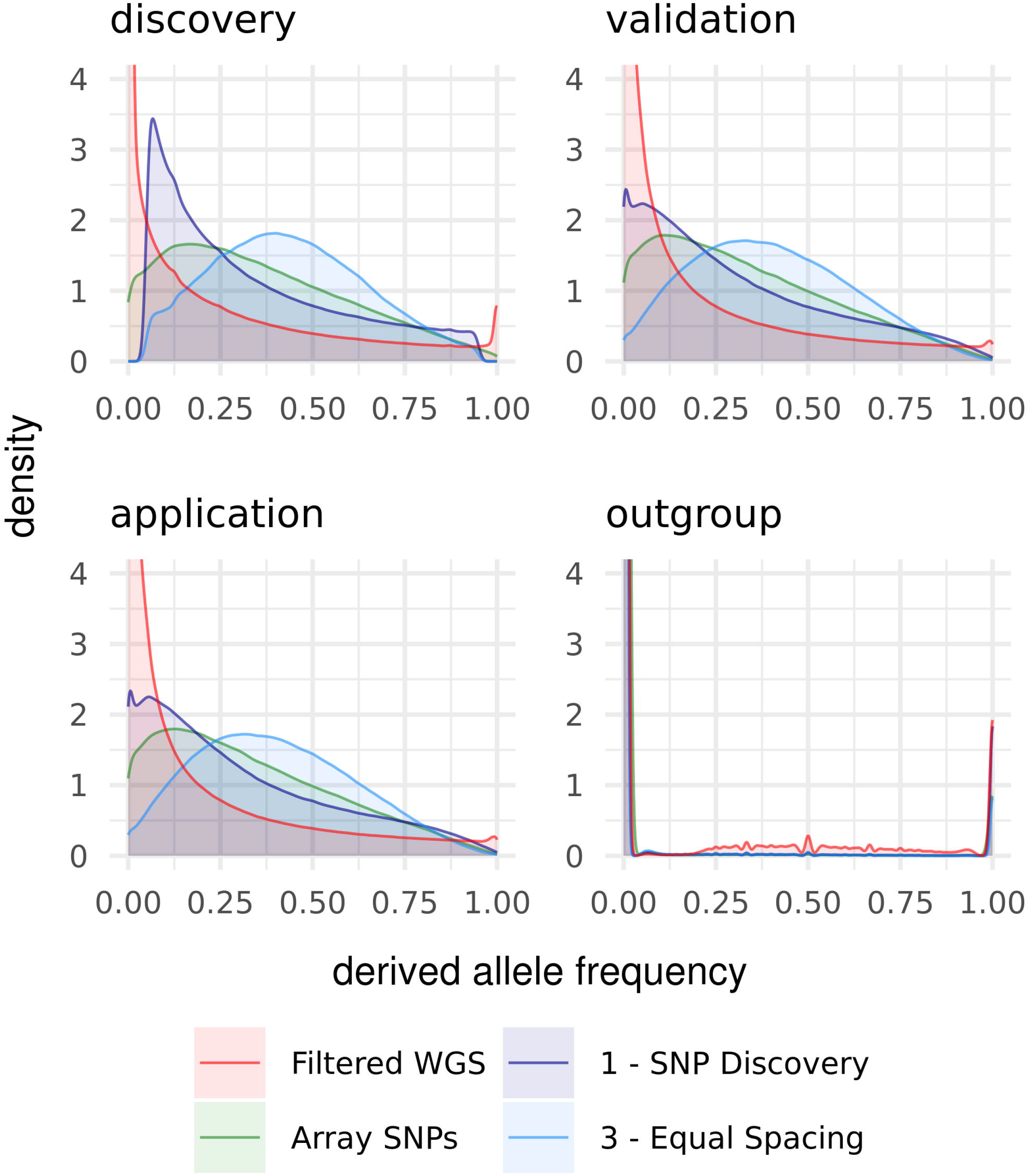
Derived allele frequency spectra within the population groups.

### Influence of number of discovery populations and target density on allele frequency spectra

Not surprisingly, an increased number of discovery populations resulted in a higher number of rare alleles after the discovery step, and thus an allele frequency spectrum with a more pronounced peak of rare alleles **(Fig 4)**. Apparently, the shift of the allele frequency spectrum after the equal spacing step was dependent on the number of discovery populations, as an increase in the number of discovery populations shifted the allele frequency spectra towards a higher proportion of alleles with a low derived allele frequency. With an increasing number of discovery populations, the shape of the allele frequency spectra got closer to the spectrum of the original array.

**Fig 4.**
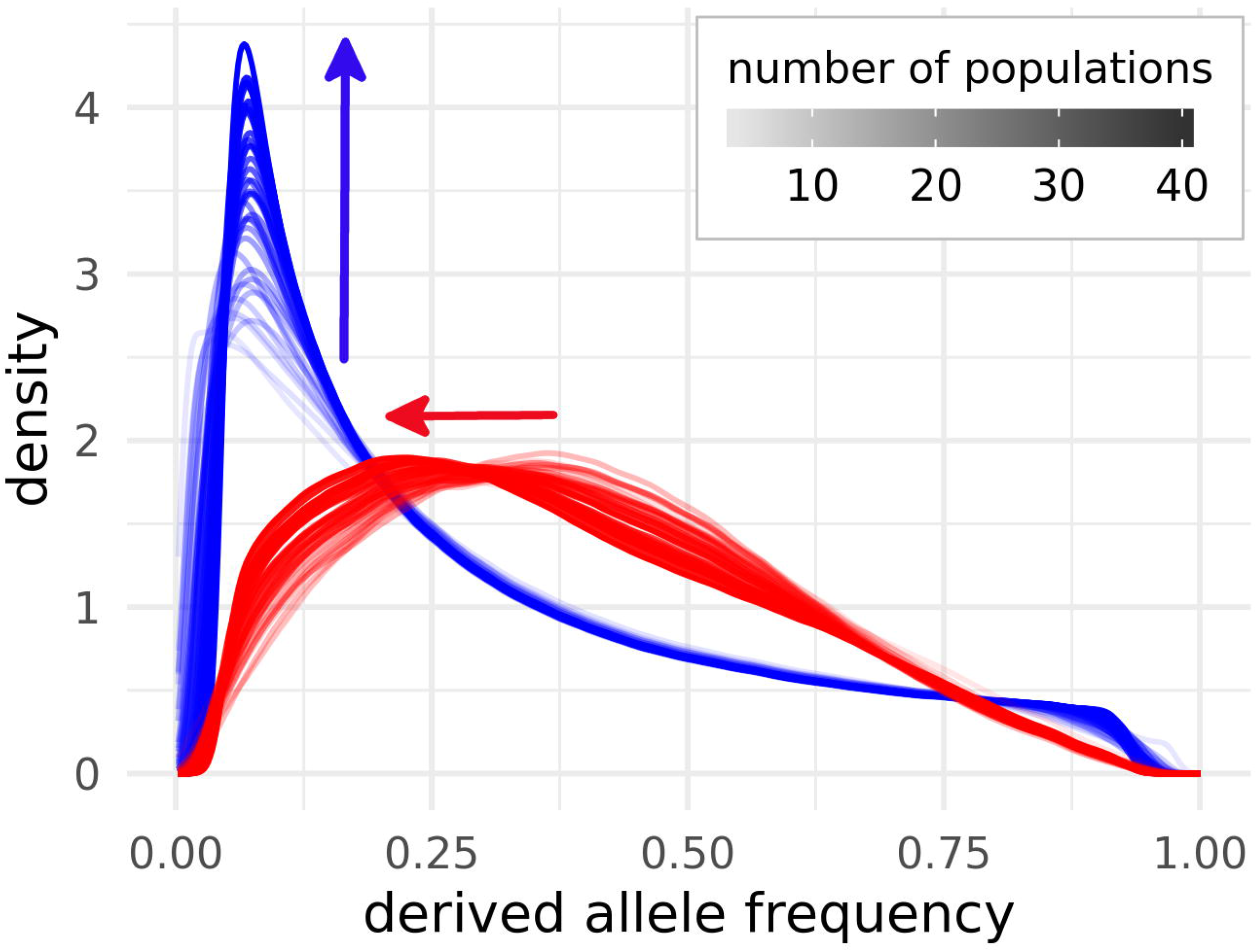
Derived allele frequency spectra after SNP discovery (blue) and equal spacing (red) for varying numbers of populations in the discovery set. Different numbers of populations in the discovery set (4 to 40) are indicated by an intensifying color gradient and only one randomly picked run per population number is shown. As the differences in the color gradients are hard to distinguish, arrows in the respective color are indicating the shift of the spectra with increasing numbers of discovery populations.

A very low target density, indicating that SNPs were mostly called due to being common backbone SNPs, resulted in an allele frequency spectrum with the majority of alleles having a MAF of around 0.5 **(Fig 5)**. Increasing the target density for the equal spacing and thus reducing the influence of the initial backbone of common SNPs shifted the peak of the allele frequency spectrum left towards a higher proportion of alleles with small derived allele frequencies. Using only the backbone SNPs common over all discovery populations and thus calling SNPs mostly by the equal spacing procedure resulted, independently from the target density, in a spectrum similar to the one obtained with a high target density with backbone **(Fig 5)**.

**Fig 5.**
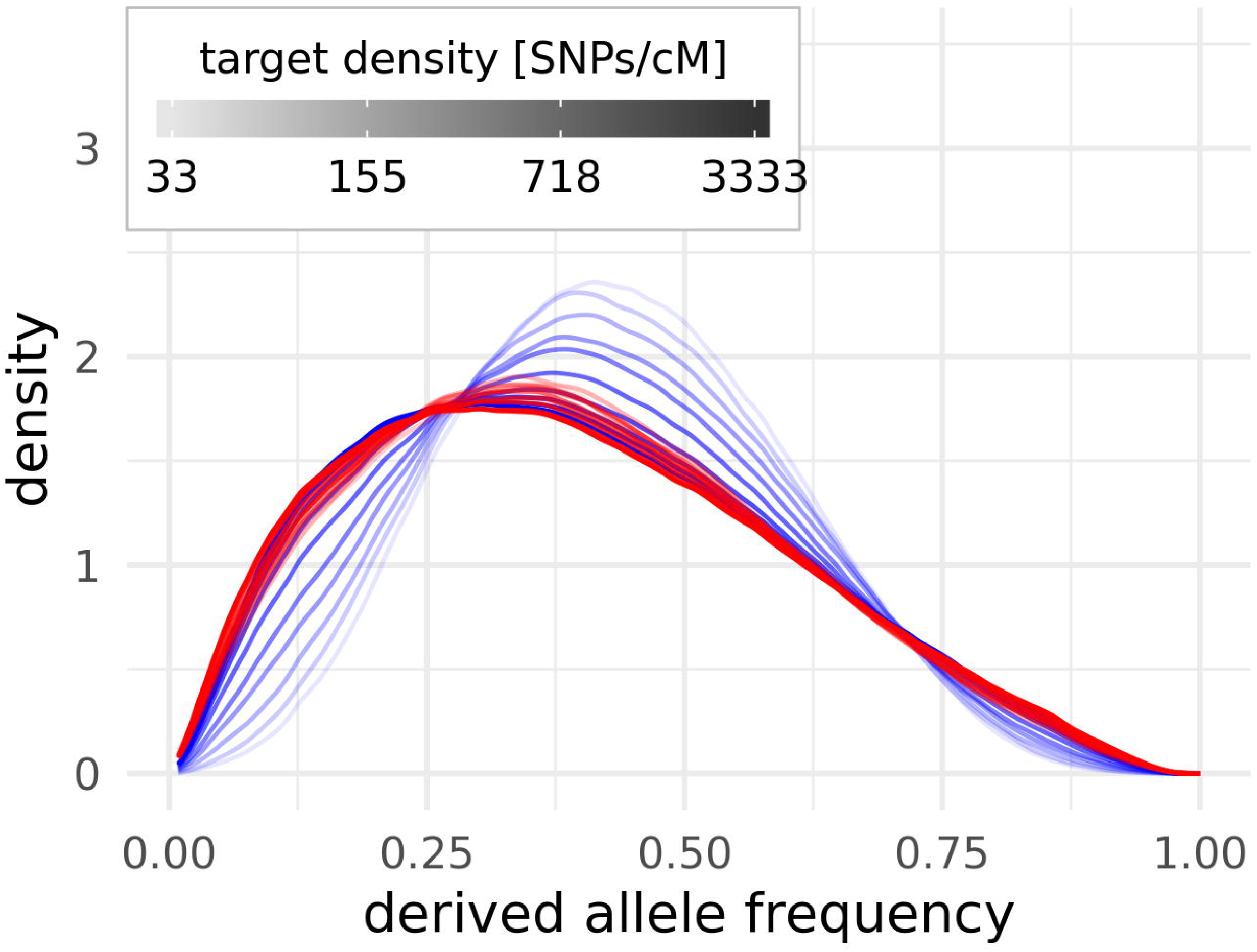
Derived allele frequency spectra for different target densities [SNPs/cM] after the equal spacing step. Blue spectra show the runs with the initial backbone included, red ones without backbone. Different target densities are indicated by a log scaled increase in color intensity. Only one representative run per target density is shown.

### Overestimation of H_exp_

**Fig 6** shows the H_exp_ of different SNP sets by population. The H_exp_ obtained from the filtered WGS SNPs were slightly higher than from the unfiltered WGS SNPs. H_exp_ obtained from the ascertained SNP sets showed an even more pronounced overestimation together with an increase during the design steps. In general, the correlations between the H_exp_ obtained in the different SNP sets were relatively high (≥ 0.95; **S2 Table)**. Especially the H_exp_ of the two WGS SNP sets showed a nearly perfect correlation of > 0.99, which led to an almost constant OHE of −0.23 **(Table 1**) for the unfiltered WGS SNPs. As already recognizable from the H_exp_ themselves, the OHE was positive for all ascertained SNP sets (0.66 – 1.29), which at the same time showed a slightly reduced correlation to the filtered WGS SNP set (0.95 – 0.97). Comparable to the allele frequency spectra, the most pronounced increase of the OHE was caused by the SNP discovery and followed by the equal spacing step (OHE increased by 0.66), while the OHE from the original array SNPs (1.41; **Fig 6; Table 1**) laid in the range covered by the remodeling steps.

**Fig 6.**
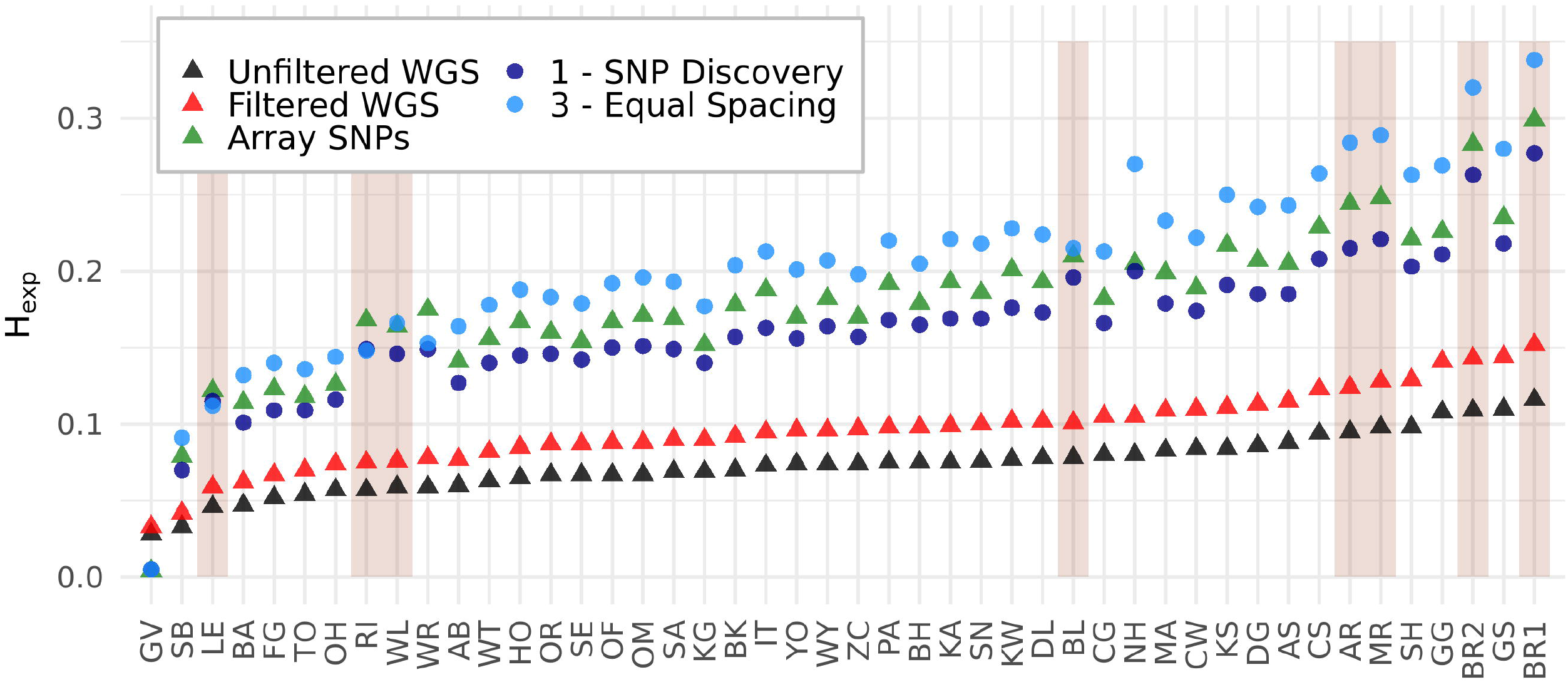
Expected Heterozygosity (H_exp_) by population and SNP set. Populations are ordered by the H_exp_ of the unfiltered WGS SNP set. Only the reference sets and relevant steps of the array design are shown. Discovery populations are shaded with a darker background.

**Table 1.** OHE of the SNP sets from the first run

| Populations | Unfiltered WGS | Array SNPs | 1 – SNP discovery | 2 - cluster removal | 3 – equal spacing | 4 – validation | 5 – down-sampling |
| --- | --- | --- | --- | --- | --- | --- | --- |
| All | -0.23 ± 0.01 | 0.84 ± 0.30 | 0.66 ± 0.26 | 0.66 ± 0.26 | 1.09 ± 0.32 | 1.29 ± 0.35 | 1.27 ± 0.34 |
| Discovery | -0.23 ± 0.00 | 1.05 ± 0.10 | 0.86 ± 0.10 | 0.86 ± 0.10 | 1.15 ± 0.15 | 1.28 ± 0.17 | 1.32 ± 0.13 |
| Validation | -0.23 ± 0.00 | 0.87 ± 0.13 | 0.68 ± 0.10 | 0.67 ± 0.10 | 1.15 ± 0.13 | 1.36 ± 0.14 | 1.33 ± 0.12 |
| Application | -0.23 ± 0.00 | 0.83 ± 0.12 | 0.64 ± 0.07 | 0.63 ± 0.07 | 1.10 ± 0.11 | 1.33 ± 0.14 | 1.30 ± 0.15 |
| Outgroup | -0.17 | -0.88 | -0.85 | -0.86 | -0.85 | -0.85 | -0.84 |

Mean OHE ± standard deviation.
An OHE of zero means no bias and an OHE of 1 means doubling the $H_{exp}$ .

Averaging the OHE within the population groups revealed a 30 % higher OHE of the discovery populations compared to validation and application populations after the discovery step. The equal spacing step reduced this difference to an only 1 % larger OHE for discovery populations, while it came with a substantial increase of the variance of OHE, which was larger for the discovery populations than validation and application populations. The validation step then increased the OHE of the validation populations more than the OHE of discovery and application populations. This stronger OHE of discovery populations was also apparent within the array SNPs (24% higher). In contrast to the other populations, the outgroup showed an underestimation of the H_exp_, resulting in an OHE of < −0.84 for all ascertained SNP sets **(Fig 6; Table 1**).

A closer look on the contribution of the sub-steps during the equal spacing step revealed that 62 % of the SNPs which were preserved during equal spacing were variable in all of the four closely related broiler populations (BR1, BR2, MR, AR; maximum pairwise Nei’s distance of 0.06 and FST of 0.17 in the filtered SNP set), while only 3 % of the SNPs were retained due to being variable in all of the four less closely related layer populations (WL, LE, BL, RI; maximum pairwise Nei’s distance of 0.15 and FST of 0.48 in the filtered SNP set). The first population used to fill in the gaps in the backbone (WL) contributed 17 % of the SNPs, while the other populations contributed < 8 %.

These findings were supported by the 50 random groupings **(Fig 7)**. The LSMEANS **(Table 2)** of the population groups revealed 24 % larger OHE for discovery populations than for validation and application populations after discovery and cluster removal step, which was decreased to a numerically insignificant difference after the equal spacing step. Interestingly, and in contrast to the findings from the first remodeling, SNP validation led to a significantly higher OHE (5 % larger) for application populations than discovery and validation populations.

**Fig 7.**
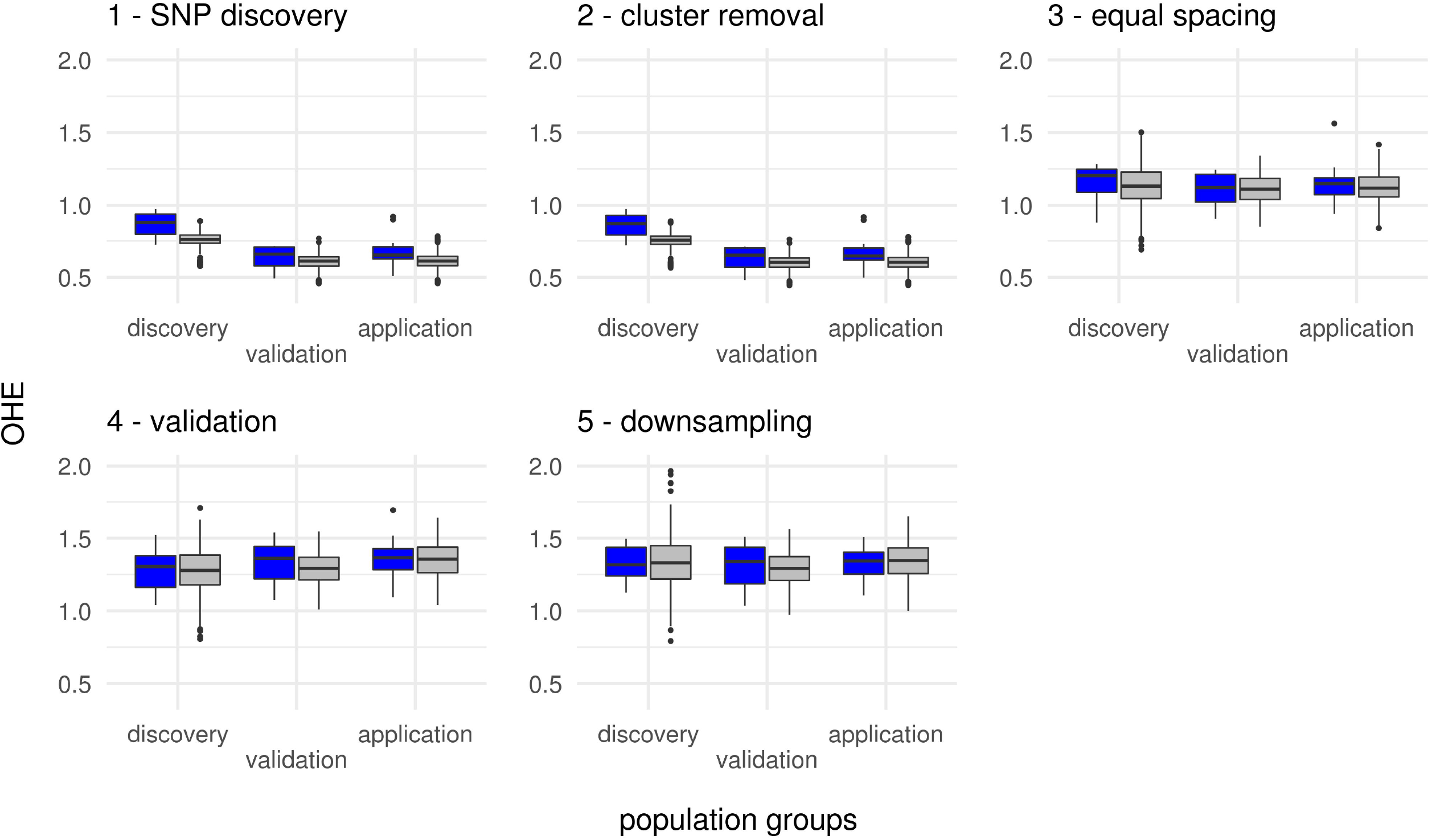
OHE for discovery validation and application populations after the five steps of array design. Discovery populations are chosen to represent populations which are comparable to the original array (blue) or 50 times random sampled (grey).

**Table 2.** OHE of the SNP sets out of the 50 random population groupings

| Populations | 1 – SNP<br>discovery | 2 – cluster<br>removal | 3 – equal<br>spacing | 4 – validation | 5 – down-<br>sampling |
| --- | --- | --- | --- | --- | --- |
| Discovery | $0.76 \pm 0.004^a$ | $0.75 \pm 0.004^a$ | $1.13 \pm 0.006^a$ | $1.28 \pm 0.006^b$ | $1.33 \pm 0.007^a$ |
| Validation | $0.61 \pm 0.003^b$ | $0.60 \pm 0.003^b$ | $1.11 \pm 0.004^b$ | $1.29 \pm 0.004^b$ | $1.29 \pm 0.005^b$ |
| Application | $0.61 \pm 0.003^b$ | $0.61 \pm 0.003^b$ | $1.12 \pm 0.004^{ab}$ | $1.35 \pm 0.004^a$ | $1.34 \pm 0.004^a$ |
| Outgroup | $-0.85 \pm 0.008^c$ | $-0.86 \pm 0.008^c$ | $-0.85 \pm 0.015^c$ | $-0.84 \pm 0.017^c$ | $-0.84 \pm 0.019^c$ |

LSMEANS for OHE ± standard error.
An OHE of zero means no bias and an OHE of 1 means doubling the $H_{exp}$ .
Different lowercase letters within columns indicate significant differences to the 5 % level.

### Influence of number of discovery populations and target density on H_exp_

**Fig 8 A** shows that increasing the number of discovery populations reduces the mean OHE of discovery populations after SNP discovery while not affecting the OHE of validation and application populations. After equal spacing **(Fig 8 B)**, no dependency of OHE on the number of discovery populations remained. As, due to the limited number of populations in the complete set, the number of validation populations decreased with more populations in the discovery set, the individual validation populations gained more impact on the ascertainment. This led to a higher OHE of validation populations with a high number of discovery populations **(S7 D Fig)**, comparable to the higher OHE of discovery populations for a small number of discovery populations.

**Fig 8.**
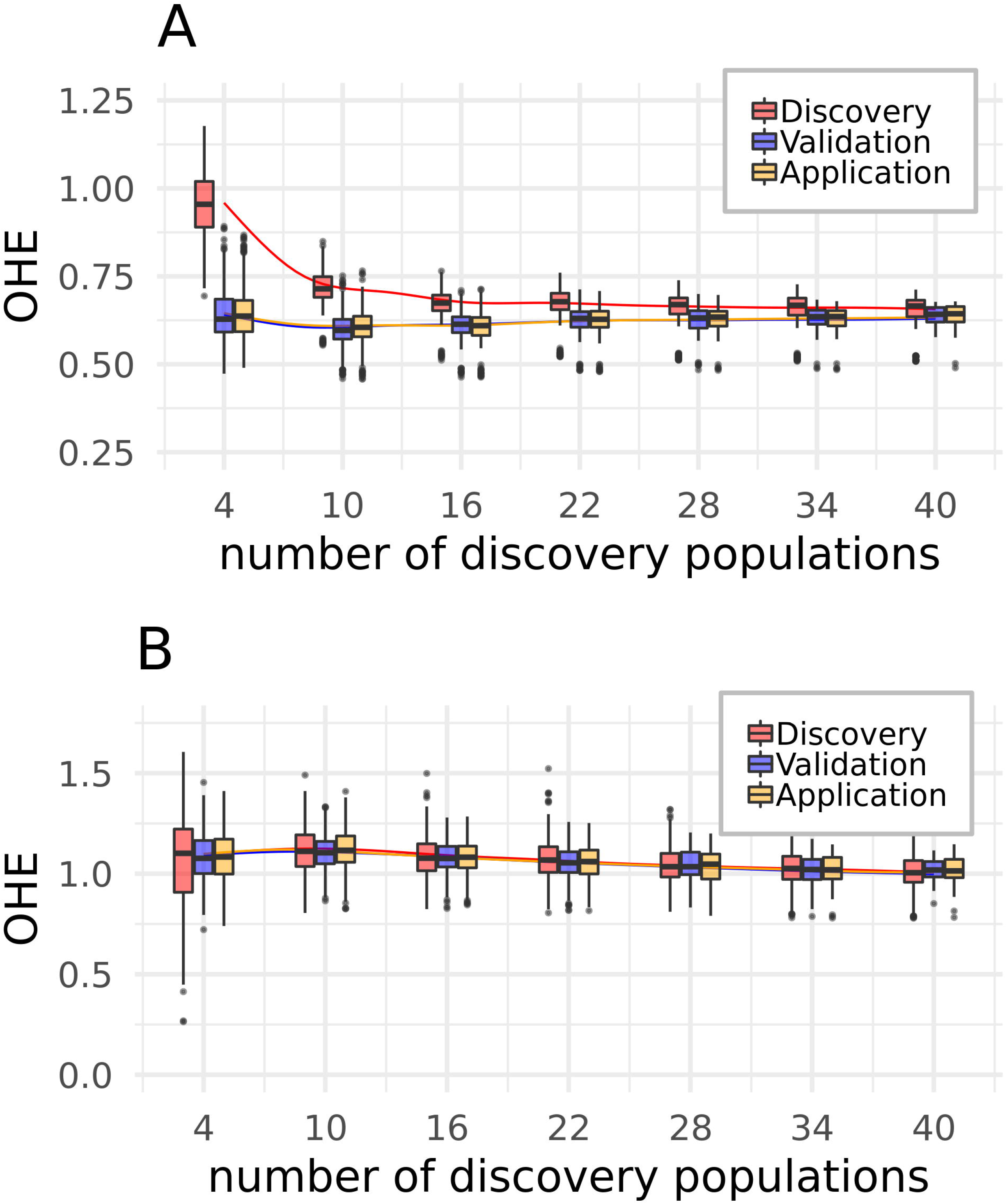
Relation of the OHE as a function of the number of discovery populations. **A -** discovery, **B -** equal spacing. The Boxplots are only shown for a subset of the number of discovery populations, while the smoothing lines, which show the trend, are calculated from all observations. Plots for all five steps can be found in **S7 Fig**.

In the equal spacing step, using only backbone SNPs resulted in a higher OHE for discovery than for non-discovery populations. Increasing the target density and thus increasing the proportion of SNPs due to the equal spacing part of the algorithm reduced the difference in OHE between the population groups **(Fig 9 A)**. If the SNPs from the initial backbone were not used, no difference of OHE between discovery and non-discovery populations was present, regardless of the target density **(Fig 9 B)**.

**Fig 9.**
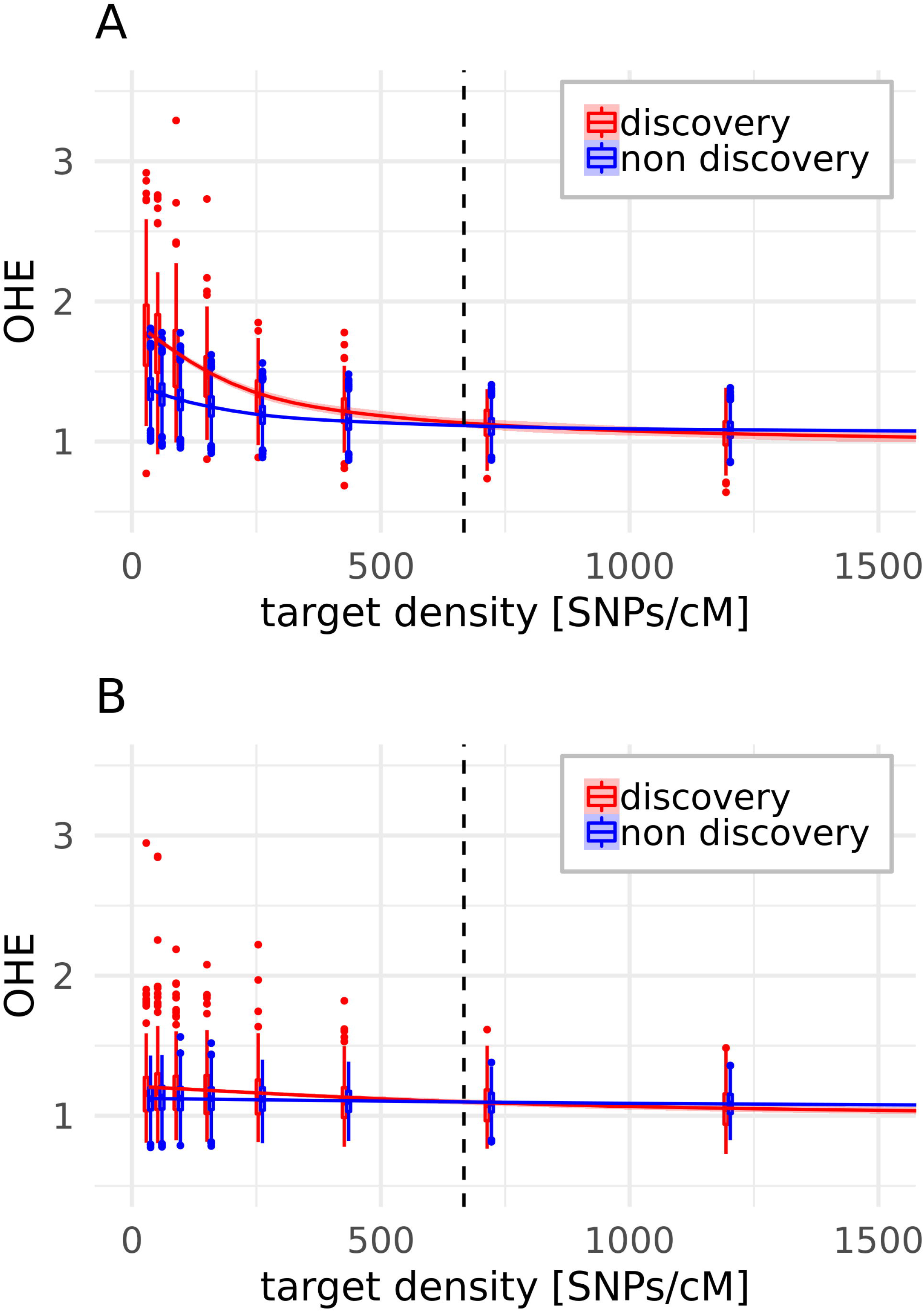
OHE after equal spacing (step 3) by target density in SNPs/cM and population group. The smoothing lines show the trend and the dashed lines the target density of 667 SNPs/cM, used for the remodeling according to the original array [21]. The algorithm was run including the initial backbone SNPs (A) or not including them **(B)**. *Gallus varius* is not included, as it is constantly underestimated.

## Discussion

In this study we used a uniquely diverse collection of sequenced wild, commercial and non-commercial chicken populations, mainly based on samples of the Synbreed Chicken Diversity Panel [13]. Parts of our set were also involved in the development process of the Axiom™ Genome-Wide 580 k Chicken Array [21]. This offered an excellent possibility for assessing the impact of ascertainment bias on real data in a complex scenario. In general, results derived in this study should therefore be comparable to other species. However, domestic chickens show a rich history of hybridization and crossbreeding events [13]. The effects of using a discovery set closely related to the commercial populations and distributing the discovery set randomly across the spectrum of populations were therefore comparably small in this study. Special patterns of population structure e.g. the stronger differentiation in cattle due to the two subspecies *Bos taurus* and *Bos indicus* [60] accompanied by limiting the discovery set to one of the two clades, should increase the impact of population structure dependent ascertainment bias.

### Potential impacts of the SNP calling pipeline

As the state of the art pipeline of GATK relies on a supervised machine learning approach for filtering the SNP calls, which needs a highly reliable set of known SNPs, we started with examining potential impacts of the filtering procedure on ascertainment bias. The number of rare variants was slightly reduced by the filtering procedure and thus increased estimates of H_exp_ were obtained in the filtered WGS set. As rare variants have a higher risk to be discarded as sequencing errors [61], this reduction is expected when applying quality filters. However, a clear assessment of correctly and falsely filtered variants is not possible here and one has to balance this tradeoff based on the study purpose.

Another source of ascertainment bias could be the use of array SNPs as training set for GATK RecalibrateVariants, which potentially leads to discarding rare variants more likely if they are not present in the discovery populations of the used array. As the correlation between the H_exp_ of the unfiltered and filtered WGS SNPs was nearly one, this source seems to be negligible and the use of array variants as a highly reliable training set seems to be unproblematic.

Due to computational limitations, we had to assume a ploidy of two for pools during the SNP calling process, which resulted in a minimal reduction of rare alleles. However, this effect was shown to have a very minor impact on the findings of this study **(S2 File)**.

### General impact over all groups

The general reduction of rare alleles in array data compared to WGS data and the resulting overestimation of H_exp_ supports findings of previous studies [25, 26, 30, 34]. This reduction of rare alleles was mainly seen at steps where selection was explicitly biased towards high MAF alleles (MAF filter for quality control in discovery step and use of common alleles for the backbone in the equal spacing step) and/ or was applied to a small number of populations (small discovery set vs. small validation set). Thereby, the strongest shifts of the allele frequency spectra and increases of H_exp_ are observed after SNP discovery and equal spacing. Both, cluster removal and second downsampling had almost no effect on the allele frequency spectra and H_exp_, while the validation step slightly decreased the share of rare SNPs.

The discovery step had the strongest impact on discovery populations, when a small set of discovery populations was used **(Fig 8 A)**. Similarly, the influence of the validation step on validation populations was strongest in case of a small number of validation populations **(S7D Fig)**. A balancing of these two groups of samples is therefore necessary, if the number of available DNA samples for array development is limited. We rather suggest the use of an extended discovery set for validation as best practice instead of using completely different populations.

If the equal spacing step contains a preselection of SNPs based on their variability within population groups, the bias is stronger towards high MAF SNPs and thus yields a higher OHE. This effect was reduced by increasing the target density and thus selecting relatively more SNPs due to the equal spacing instead of common occurrence.

### Differences between groups

If allele frequency spectra are changed in the same way for all populations and are therefore biasing heterozygosity estimates to the same extent, findings for between population comparisons will be little affected. Ascertainment bias then is only of importance if one compares populations based on different arrays, and corrections of the allele frequency spectrum as reviewed by Nielsen [25] should be possible. As correlations between H_exp_ of ascertained SNP sets and unfiltered/ filtered WGS SNP sets were consistently high (> 0.94), arrays designed in the way as performed in this paper should mostly be suitable for robust and cost efficient analyses. Biasedness of estimates could be reduced even more by considering filter strategies according to Malomane *et al*. [30].

However, we could show that the bias acts with different extent on different population groups (population structure dependent bias) and therefore changes ranking of populations and can affect conclusions. This population structure dependent bias was already shown to have severe impact on findings from SNP arrays. For example, Bradbury *et al*. [62] found a demographic decline up to an approximately 30% lower H_exp_ for Atlantic cod based on the distance to the sampling location of the discovery panel and McTravish and Hillis [29] showed strong deviations between simulated and observed polymorphisms for different combinations of migration- and ascertainment scenarios on simulated cattle populations. In concordance with this, populations which are closely related to the discovery populations of the original array in our study on average showed a 24 % higher OHE than validation and application populations for the original array.

This population structure dependent bias was mainly introduced by the initial discovery step. It was also observed for the random population groupings, but to a slightly different extent. The difference in overestimation decreased with an increase in the number of discovery populations **(Fig 8)** and was smallest if the discovery populations showed minimum distance to the application and validation populations (results not shown). Comparable observations were already made by Frascaroli *et al*. [63] which found very small ascertainment bias for European elite maize lines when using a SNP panel discovered in a combination of a maize diversity set and inbred lines, but strong ascertainment bias when using SNPs which were discovered in American elite lines. Therefore, we suggest to ideally choose an array where the discovery panel does span the scope of populations it will be applied to, and by this covers the existing variation in a most representative way, or to design such an array for oneself if it does not exist.

The equal spacing step lowered the difference in mean OHE between population groups in most of our remodeling scenarios, but obviously not in case of the original array. In the remodeling, we saw this difference only with a low target density and thus calling SNPs in the equal spacing step mainly due to being common over many populations **(Fig 9 A)**. However, the equal spacing step also increased the variance of OHE in the discovery panel, meaning that the OHE was increased more for some of the discovery populations than for others, thus causing more uncertainty for resulting effects. This effect is driven by the unequal contribution of variable SNPs to the chosen SNP set by the different populations during the equal spacing step **(S3 Fig)**. The equal spacing step increases the OHE for some of the discovery populations, while it decreases it for others, and hence it does not remove the population structure dependent bias. This means that the knowledge of which discovery populations were used is not sufficient to draw conclusions regarding a possible ascertainment bias, since their relative contribution varies through the described pipeline.

### Outgroup

*Gallus varius* as an outgroup showed a different behavior than all other populations. It already exhibited the lowest H_exp_ in the unfiltered WGS SNP set, which was most likely driven by the small number of only two samples in the pool, and showed less upward bias of H_exp_ in the filtered WGS SNP set than all other populations. The *Gallus varius* sequence reads on average showed weak Phred-scaled mapping quality scores of 19 (1.3 % probability of misalignment), while the mean quality scores of the other populations ranged from 25 (0.3 %) to 28 (0.1 %) – meaning that variation, only present in *Gallus varius*, will be more likely missed due to misplacement of the reads or discarded as possible sequencing errors. Additionally, every ascertained SNP set showed an OHE for *Gallus varius* of < −0.84, as variation being present only in *Gallus varius* was not found in *Gallus gallus* discovery panels and contrary variants from *Gallus gallus* were not variable in *Gallus varius* **(Fig 3)**. This demonstrates that arrays should not be used if different species (even closely related ones) are included in the research project. Even sequence based estimates can be slightly biased, if the reference genome does not fit properly.

### Potential impact on other breeding applications

In general, we cannot infer the impact on breeding applications which require phenotypic data (e.g. genomic selection [64] or genome wide association studies [65]) and/or individually sequenced or genotyped individuals (e.g. linkage disequilibrium decay [66] or runs of homozygosity analyses [67]) from this study. However, literature highlights the increased power of high MAF SNPs to capture/ detect effects which are caused by common variants due to stronger linkage disequilibrium and higher levels of variance explained. Therefore, increasing MAF in a first instance increases prediction accuracy when the number of SNPs is limited [68] and therefore some SNP ascertainment schemes intentionally bias the used SNPs towards high MAF within the desired populations [17]. The switch to WGS data, and therefore the additional inclusion of rare alleles, is then expected to increase the possibility of capturing the effects of rare alleles [68–70]. However, the increase in efficiency by higher numbers of SNPs levels off when going towards WGS data [71]. Nevertheless, we would expect negative impacts of ascertainment bias due to the across population use of the arrays. When biasing the genotyped variation towards the discovery population, the variability in populations, which are less related to the discovery populations, is less increased or even reduced, and arrays therefore become less valuable in non-target populations. Slight effects of this were demonstrated by simulation [68] and we can clearly support these findings by the levels of differences in the genotyped heterozygosity which we observed in this study. For the effect of ascertainment bias on a broader set of applications, we further refer the interested reader to studies which specifically address those issues (e.g. 30,31,66).

### Further recommendations for future studies

We showed that existing arrays come with a large, and due to a diverse set of promoting factors barely predictable, potential for ascertainment bias. Strongly decreasing costs for WGS and increasing availability of powerful computing resources therefore promote an intensified use of WGS for population genetic analyses, especially when diverse populations are included in the studies. However, costs and computational effort will still be substantial for large scale projects. Possible cost effective alternatives could be reduced library sequencing approaches like Genotyping-by-Sequencing [61, 72], even though such methods introduce other problems related to the use of restriction enzymes which are reviewed by Andrews *et al*. [73].

For the purpose of monitoring genetic diversity in a large set of small non-commercial populations, the development of a specialized new array for cost effective high throughput genotyping could be still a good option. For the design of such an array, it is crucial to extend the discovery set in a way which represents the total variability over all populations as balanced as possible. The use of publicly available sequences can be helpful to reach this goal. The ascertainment of the SNPs should then be done preferably over a large set of highly diverse populations covering a wide spectrum of the diversity within a species available populations instead of biasing the process towards common alleles by performing within population ascertainment.

However, when considering the findings of this study, choosing an array for which the SNP ascertainment process is well documented and a discovery panel that matches the populations to be studied, should enable researchers to derive appropriate conclusions when robust estimators are used.

## Acknowledgments

We are grateful to all the participating breeders for their assistance in sampling.

## Supporting information

**S1 File. Abbreviations for breeds and accession numbers for the sequencing data**.

**S2 File. Supplementary methods**.

**S3 File. Pairwise Nei’s standard genetic distances**.

**S4 File. Pairwise F_ST_ values**.

**S1 Fig. Pooled vs. ploidy two calling**. Expected Heterozygosity (H_exp_) from the calling with assuming the correct ploidy vs. assuming ploidy two for all samples (A) and alternative allele frequency distributions of called alleles for individually sequenced (B) and pooled sequenced (C) populations.

**S2 Fig. Alternative allele frequency spectrum for SNPs where reference allele is not ancestral allele (A) and alternative (B) vs. derived (C) allele frequency spectrum of all SNPs**.

**S3 Fig. Cumulative number of SNPs in million retained during the equal spacing step**. The red line and points represent the first remodeling according to the original array [21], while the dashed lines and the boxplots represent the 50 random population groupings and the black line the according median values. The algorithm starts with a very basic initial back-bone and then adds SNPs to the backbone which are variable in either all layer lines or all broiler lines. Separated by a vertical line, the second part of the algorithm successively fills up potential gaps to achieve an equidistant coverage of 667 segregating SNPs/cM for each discovery population.

**S4 Fig. SNPs retained by number of populations in the discovery set**.

**S5 Fig. Number of SNPs retained after the equal spacing step by target density**.

**S6 Fig. Relative amount of SNPs shared by a specific number of runs from 50 random population assignments**.

**S7 Fig. Relation of the OHE as a function of the number of discovery populations**. **A** - discovery, **B** – cluster removal, **C** – equal spacing, **D** – validation, **E** – downsampling. The Boxplots are only shown for a subset of the number of discovery populations, while the smoothing lines, which show the trend, are calculated from all observations.

**S1 Table. Number of SNPs from the remodeling processes**

**S2 Table. Pearson correlations between the H_exp_ of the different SNP sets**

