## SupplementaryMaterial for "How Array Design affects SNP Ascertainment Bias": S1_Table.docx

Table S 1: Number of SNPs from the remodeling processes

|  | 1 – SNP  discovery | 2 - cluster removal | 3 – equal spacing | 4 – vali­dation | 5 – down­sampling |
| --- | --- | --- | --- | --- | --- |
| Original Array (Kranis *et al.* 2013) | 23 998 | 10 029 | 1 829 | 1 187 | 581 |
| First Run | 10 903 | 8 836 | 2 050 | 1 714 | 580 |
| Repeated Remodeling | 11 603 ± 226 | 9 328 ± 157 | 2 032 ± 147 | 1 698 ± 147 | 662 ± 33 |
| Number of SNPs in 1,000  In case of repeated remodeling mean of the 50 runs ± standard deviation | | | | | |
