## SupplementaryMaterial for "How Array Design affects SNP Ascertainment Bias": S2_File.docx

Supplementary file 2
- supplementary methods -

### Correct ploidy calling vs. ploidy two calling

Due to the large diversity of the chicken populations and a series of bugs in GATK (up to version 4.2), it was only possible to call some intervals, spanning 28.2 % of the autosomal region, with the correct ploidy. Compared to the calling with the correct ploidy, assuming ploidy two for the pools came along with a slight reduction of SNPs with very rare alternative allele frequencies of ≤ 0.02 in the pools (**S1C Fig**). As those SNPs had to be invariable in the individually sequenced populations, a reduction of invariable alleles is present in the individually sequenced populations (**S1B Fig**). When comparing expected heterozygosities (H_exp_) from the two callsets, the ploidy two calling comes along with a slight increase of H_exp_, which is minimally larger for individually sequenced populations (**S1A Fig**), as the lost SNPs are invariable rather than rare in those populations. However, as the correlation between the two callsets is > 0.99 and the mean overestimation of H_exp_ (OHE) for the pools is 0.02 and 0.05 for the individual sequenced populations, the effect is negligible compared to the effects of ascertainment bias.

### Alternative allele frequency spectrum vs. derived allele frequency spectrum

As the chicken reference genome is based on an inbred Red Jungle Fowl (*Gallus gallus gallus*) [1], many publications handle the reference allele as ancestral allele (e.g. [2]). However, this assumption does not account for mutation, selection and inbreeding happening in the Red Jungle Fowl after separation from the domesticated strains as well as variation already present before separation. Therefore, ancestral alleles were defined by an approach comparable to Rocha *et al.* [3] and using allele frequency information from the three wild populations *Gallus gallus gallus* (GG), *Gallus gallus spadiceus* (GS) and *Gallus varius* (GV). It was assumed that the *Gallus gallus* and *Galllus varius* species emerged from a common ancestor and *Gallus gallus* later split into *Gallus gallus gallus* and *Gallus gallus spadiceus* subspecies. Additionally, assuming neutral molecular evolution [4], the ancestral allele was most likely the major allele within those three populations, when weighting the allele frequency of *Gallus varius* twice. This procedure assigned the ancestral status to the reference allele for 86 % of the SNPs and to the alternative allele for 14 % of the SNPs. SNPs where the ancestral state was assigned to the alternative allele were approximately equal distributed over the range of allele frequencies (**S2A Fig**). Only alleles being close to fixation for the alternative allele were overrepresented, possibly being artefacts of selection on de novo mutations in ancestors of the reference individual or sequencing errors in the reference genome. Therefore, a comparison between alternative (**S2B Fig**) and derived (**S2C Fig**) allele frequency spectrum in the complete callset does only show the reduction of a tiny peak being present at the end of the alternative allele frequency spectrum (111’851 SNPs; 0.039 % of all SNPs) which goes along with an increase of the first 5 % bin of the same number. All other changes in the number of SNPs per bin were less than 0.05 % of all SNPs.
