## SupplementaryMaterial for "How Array Design affects SNP Ascertainment Bias": S2_Table.docx

Table S 2: Pearson correlations between the H_exp_ of the different SNP sets

|  | Filtered WGS | Array SNPs | 1 – SNP discovery | 2 - cluster removal | 3 – equal spacing | 4 – vali­dation | 5 – down-sampling |
| --- | --- | --- | --- | --- | --- | --- | --- |
| Unfiltered WGS | 1.00 | 0.94 | 0.95 | 0.95 | 0.96 | 0.96 | 0.96 |
| Filtered WGS |  | 0.95 | 0.96 | 0.95 | 0.97 | 0.96 | 0.96 |
| Array SNPs |  |  | 1.00 | 1.00 | 0.98 | 0.97 | 0.98 |
| 1 – SNP discovery |  |  |  | 1.00 | 0.98 | 0.97 | 0.97 |
| 2 - cluster removal |  |  |  |  | 0.98 | 0.96 | 0.97 |
| 3 – equal spacing |  |  |  |  |  | 1.00 | 0.99 |
| 4 – vali­dation |  |  |  |  |  |  | 1.00 |
