## Supplementary figures and images for "How Array Design affects SNP Ascertainment Bias"

### S1_Fig.tiff

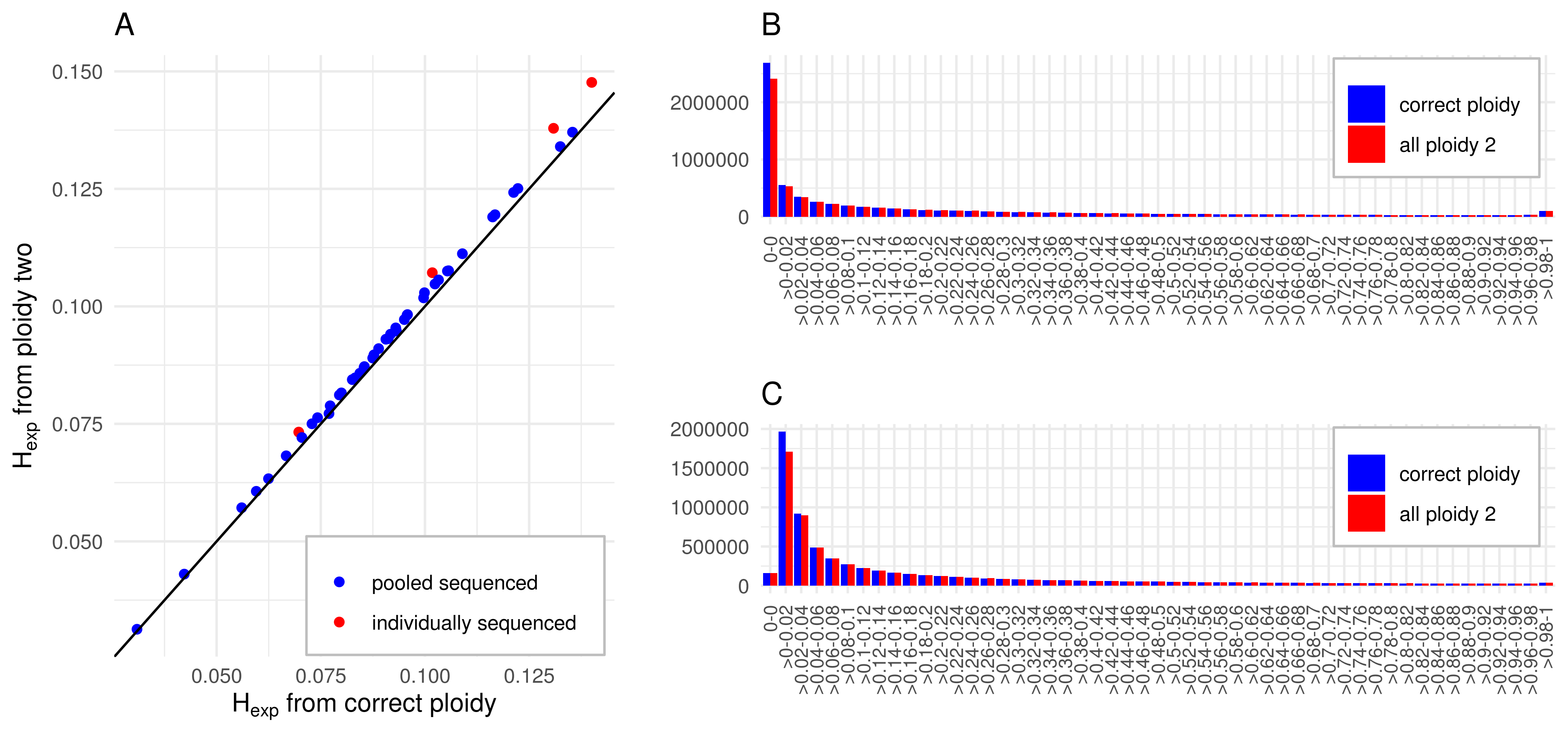

### S2_Fig.tiff

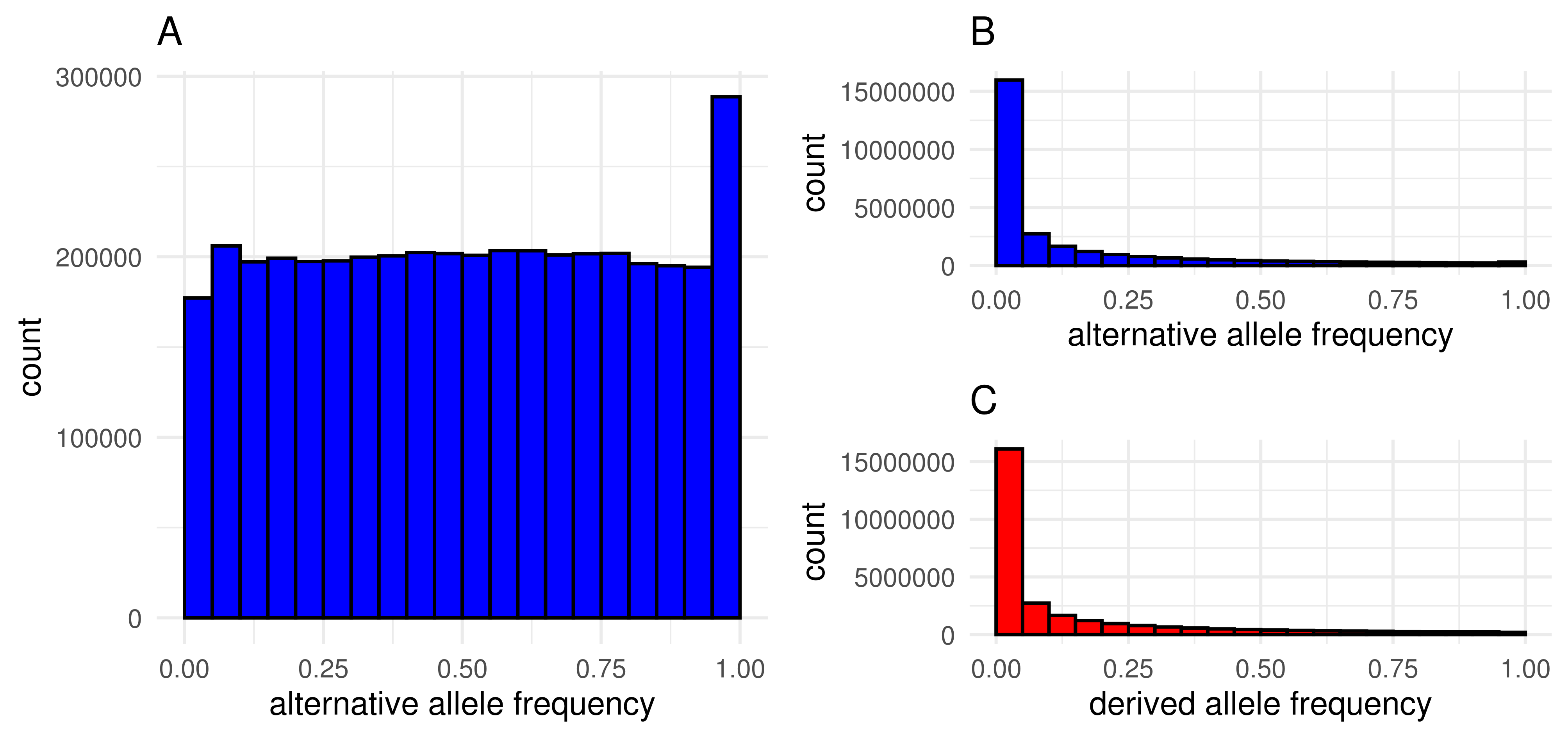

### S3_Fig.tiff

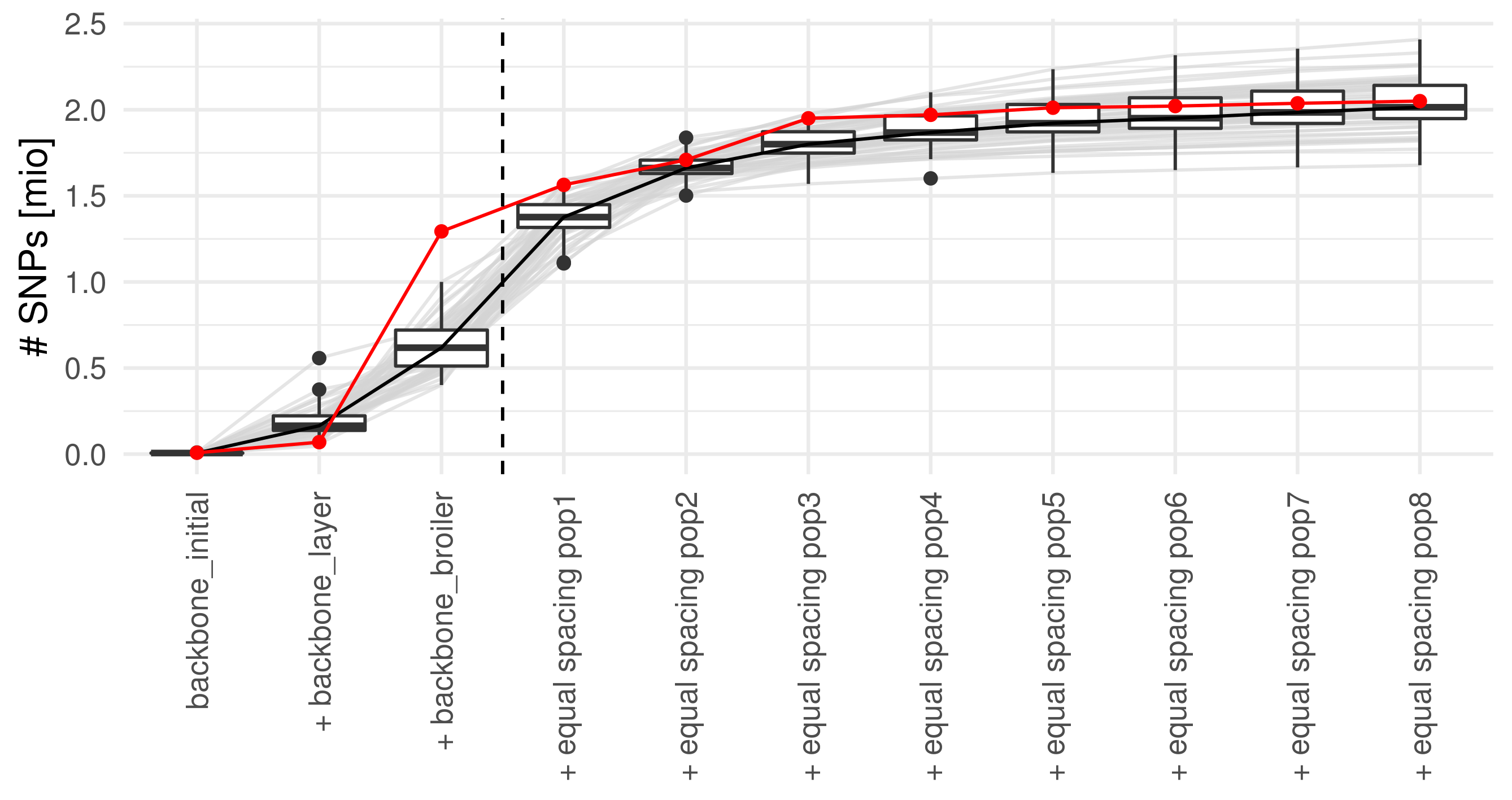

### S4_Fig.tiff

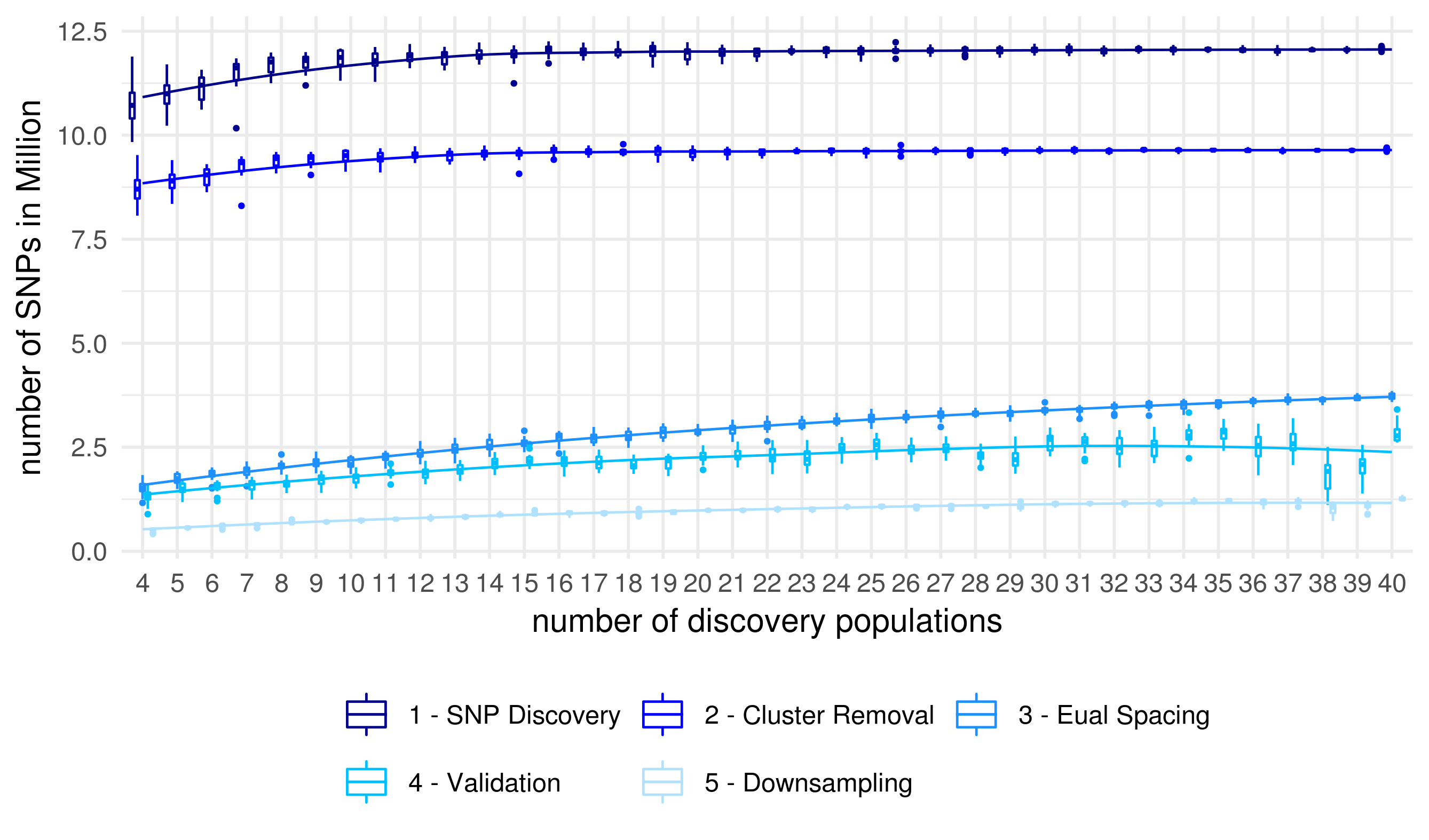

### S5_Fig.tiff

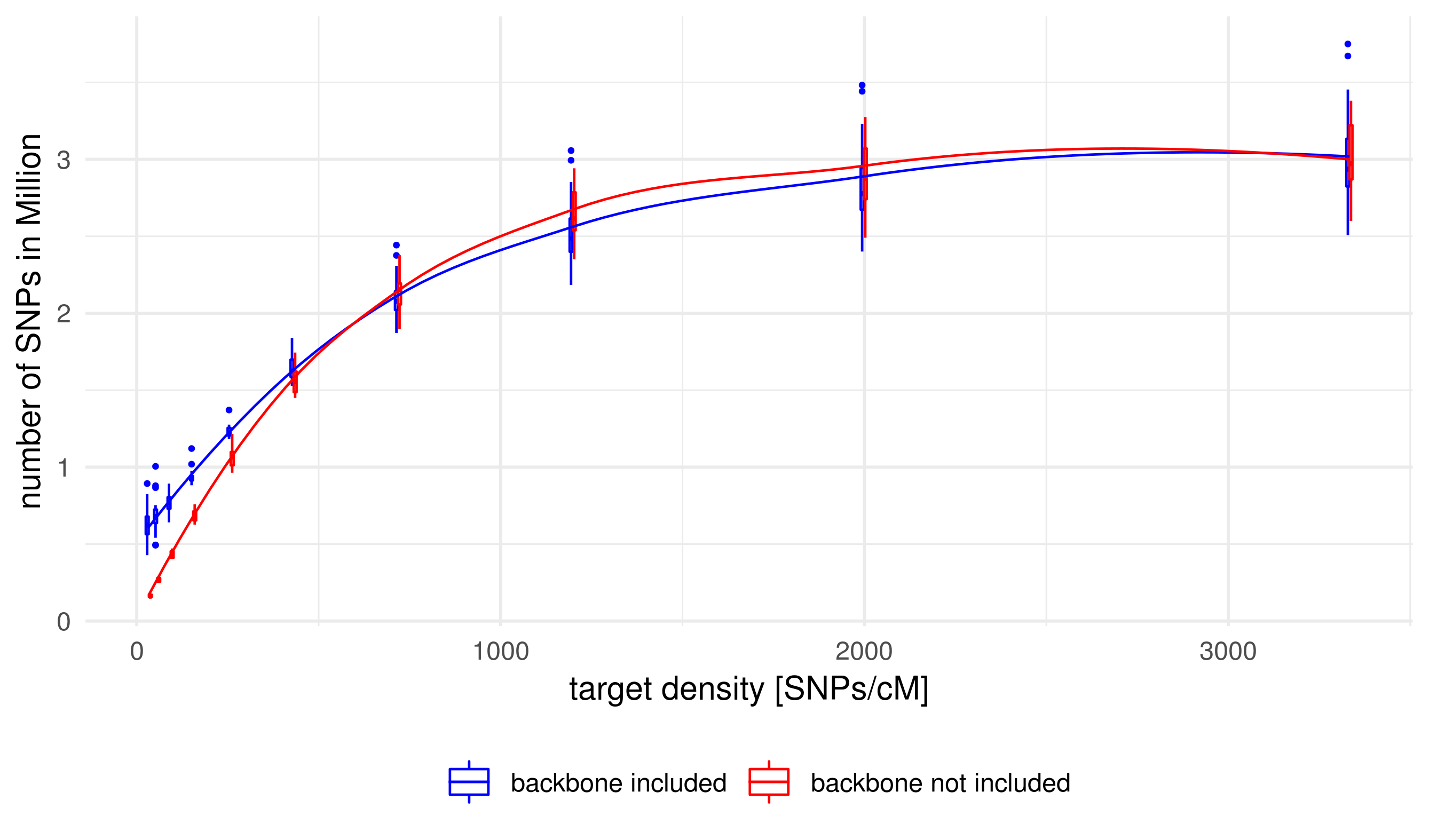

### S6_Fig.tiff

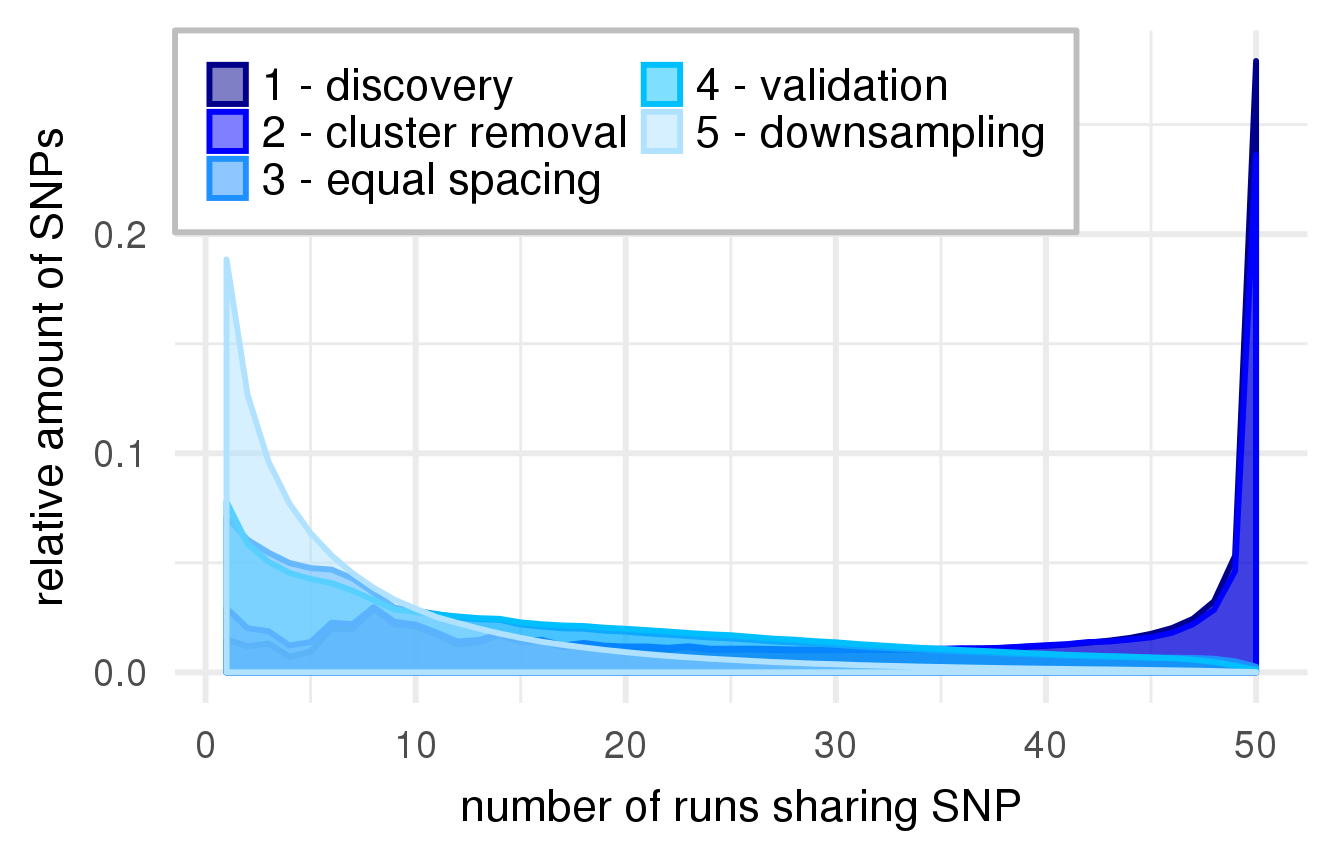

### S7_Fig.tiff

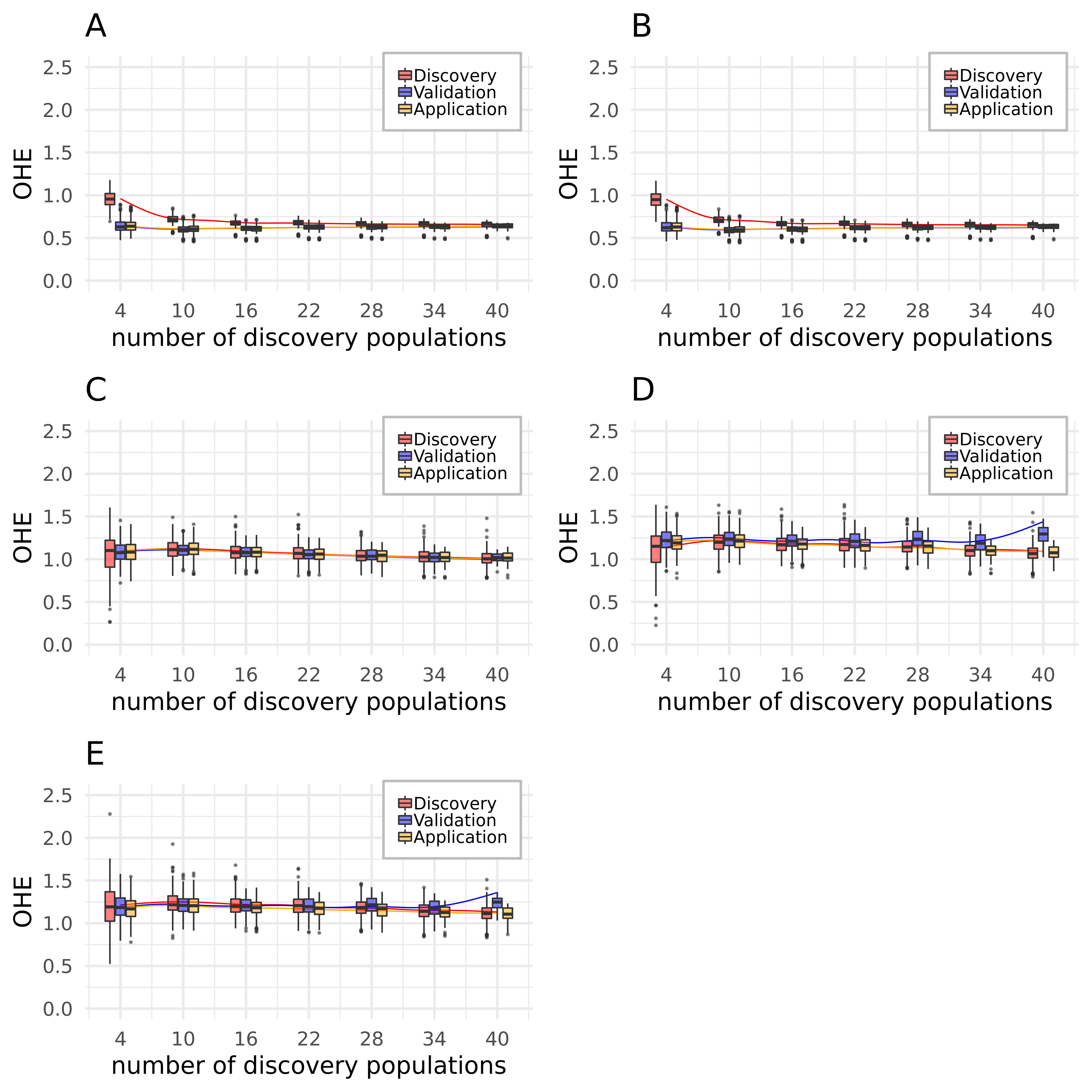
